# Human Whole Genome Sequencing in South Africa

**DOI:** 10.1101/2020.06.10.144402

**Authors:** Brigitte Glanzmann, Tracey Jooste, Samira Ghoor, Richard Gordon, Rizwana Mia, Jun Mao, Hao Li, Patrick Charls, Craig Douman, Maritha J. Kotze, Armand V. Peeters, Glaudina Loots, Monika Esser, Caroline T. Tiemessen, Robert J Wilkinson, Johan Louw, Glenda Gray, Robin M. Warren, Marlo Möller, Craig Kinnear

**Affiliations:** DSI-NRF Centre of Excellence for Biomedical Tuberculosis Research, SAMRC Centre for Tuberculosis Research, Division of Molecular Biology and Human Genetics, Faculty of Medicine and Health Sciences, Stellenbosch University, Cape Town, South Africa; Biomedical Research and Innovation Platform, South African Medical Research Council, Tygerberg, Cape Town, South Africa; Division of Medical Physiology Faculty of Medicine and Health Sciences, Stellenbosch University, Tygerberg Hospital, Cape Town, South Africa; Grants, Innovation and Product Development, South African Medical Research Council, Tygerberg, Cape Town; BGI-Shenzhen-Building 11, Beishan Industrial Zone, Yantian District, Shenzhen (518083); Information Technology Services Division, South African Medical Research Council, Cape Town, South Africa; Division of Chemical Pathology, Department of Pathology, Faculty of Medicine and Health Sciences, Stellenbosch University, Cape Town, South Africa; Division of Chemical Pathology, Department of Pathology National Health Laboratory Service, Tygerberg Hospital, Tygerberg, South Africa; South African National Department of Science and Innovation, South Africa; Department of Paediatrics and Child Health, Faculty of Medicine and Health Sciences, Stellenbosch University, Tygerberg Hospital, Cape Town, South Africa; Centre for HIV and STIs, National Institute for Communicable Diseases, and Faculty of Health Sciences, University of the Witwatersrand, Johannesburg, South Africa; Wellcome Centre for Infectious Diseases Research in Africa, Institute of Infectious Disease and Molecular Medicine, University of Cape Town, Observatory 7925, South Africa; Department of Infectious Diseases, Imperial College London, W12ONN, United Kingdom; The Francis Crick Institute, London, NW1 1AT, United Kingdom; Genomics Centre, South African Medical Research Council, Tygerberg, Cape Town, South Africa; Office of the President, South African Medical Research Council, Cape Town, South Africa; Perinatal HIV Research Unit, Faculty of Clinical Medicine, University of the Witwatersrand, Chris Hani Baragwanath Academic Hospital, Johannesburg, South Africa

## Abstract

The advent and evolution of next generation sequencing has considerably impacted genomic research. Until recently, South African researchers were unable to access affordable platforms capable of human whole genome sequencing locally and DNA samples had to be exported. Here we report the whole genome sequences of the first six human DNA samples sequenced and analysed at the South African Medical Research Council’s Genomics Centre. We demonstrate that the data obtained is of high quality, with an average sequencing depth of 36.41, and that the output is comparable to data generated internationally on a similar platform. The Genomics Centre creates an environment where African researchers are able to access world class facilities, increasing local capacity to sequence whole genomes as well as store and analyse the data.

## Introduction

The Human Genome Project (HGP) resulted in the completion of the first human genome sequence, a major breakthrough in the field that propelled genetic studies. This project depended on Sanger sequencing and took 13 years, a large international collaboration and approximately $300 million to complete ^1^. The ensuing development of next-generation sequencing (NGS) technologies made it possible to rapidly sequence large amounts of DNA at an affordable cost, which substantially impacts clinical practice, particularly clinical genetics and oncology, as well as human genetic research ^2^.

The Beijing Genomics Institute (BGI) participated in the original HGP and subsequently developed many of its own sequencing instruments. The BGISEQ-500 was the first platform capable of competing with Illumina’s instruments and offered high quality sequencing at a reduced cost ^3^ (3). MGI Tech Co.Ltd (MGI), a subsidiary of BGI, released two new sequencing instruments, namely the MGISEQ-2000 and MGISEQ-200, in October 2017. These sequencers rely on MGI’s proprietary DNBseq™ technology and the combinatorial Probe-Anchor Synthesis (cPAS) method, an improvement of the combinatorial Probe-Anchor Ligation (cPAL) sequencing technology, first patented by Complete Genomics ^4,5^. As described by Korostin et al, the MGI-SEQ2000 is a complete alternative to the Illumina platform for similar tasks, including whole genome sequencing (WGS) ^4^. Importantly, the affordable pricing has made it possible to provide human WGS in settings with limited resources, such as South Africa.

Human genetic studies in African countries hold much promise, but are more challenging to do than elsewhere, resulting in the underrepresentation of populations from this continent ^6,7^ even though African researchers have the proven capacity to conduct large-scale human genetic analyses. For example, the Southern African Human Genome Programme (SAHGP) investigated the whole genomes of 24 individuals ^8^, while the H3ABionet consortium has a node in South Africa and developed African bioinformatics infrastructure ^9^. However, because South African researchers were previously unable to access platforms capable of human WGS locally, DNA samples had to be exported. It is a legal requirement that export permits for samples must be obtained from the South African Department of Health, which can only be applied for once a service contract has been reviewed by legal advisors, signed by the representative of the research institute and submitted together with proof of ethics approval ^10,11^. In addition to this, the demand for export permits means that in some cases, researchers may wait up to a few months for an export permit, significantly impacting research timelines. In 2019, the South African Medical Research Council (SAMRC), in partnership with the BGI, launched the first high throughput WGS platform in South Africa. The local availability of WGS makes exporting of samples unnecessary and expedites human genetic research in one of the most diverse countries in the world. Additionally, it allows researchers in South Africa to produce and analyse African genomics data on African soil, at an affordable price.

Here we present the whole genome sequences of the first six human samples sequenced and analysed in South Africa at the SAMRC Genomics Centre. We further compare the results obtained from the South African installation of the MGISEQ-2000 to that of the same sample sequenced on a BGISEQ-500 at the BGI, China. Three DNA samples of known genotype previously determined by whole exome sequencing (WES) covering only the coding regions of the human genome, enabled limited analytical validation and assessment of diagnostic accuracy in a family with Li Fraumeni-like syndrome.

## Results

### Comparison of sequencing and mapping data quality

A total of six genomic DNA samples were sequenced at the SAMRC Genomics Centre, one of which was also sequenced at the Beijing Genomics Institute in China, as a means of comparison. All individual fastq files were processed identically (Supplementary Figure 1). Basic summary statistics of the data are shown in Table 1. Raw fastq sequences were analysed using FastQC ^12^ and these results illustrate that all of the outputs were of high quality (Supplementary Figures 1–7). Reads were subsequently preprocessed by trimming 5 base pairs (bp) from each end of the read to remove potential low-quality reads and possible adaptor contamination. Following individual analysis of the raw data, all fastq files with different barcodes were merged into their individual forward and reverse reads. FastQC was repeated to ensure that the data quality remained acceptable. Of the samples analysed, 91% of sequenced bases had a base quality score of more than 30. An average coverage of 36.48X was obtained for all of the samples and this coverage remained consistent across the entire read length of 100 base pairs (bp) (Figure 1). The per-sequence quality scores were consistent for all samples across the length of the reads (Figure 2) and the GC content, plotted against the theoretical GC content of the reference genome, was uniform across all seven samples (Figure 3). Trimmed and filtered reads were aligned to the human reference genome GRCh38p13 using Burrows Wheeler Aligner-MEM, and the quality of the read alignments was assessed using the bamstats module in SAMtools ^13^. Read quality was acceptable for each of the samples with the proportion of aligned reads averaging 99.57% across all samples. Average insert size for each of the libraries was 257bp (range of 251bp and 263bp respectively), as per the manufacturer’s protocol which suggests between 250bp and 300bp. Furthermore, it was determined that none of the samples had duplicate reads and BQRS was performed to ensure that mismatches in the alignment were corrected. The error rate, which is calculated as mismatches per base mapped for each of the samples, is shown in Figure 4. In total, an average of 4,695,160 variants per sample were identified in all samples sequenced, with 3,752,860 (230,432 novel) single nucleotide polymorphisms (SNPs) and 941,871 (226,151 novel) insertions/deletions (Figure 5, Supplementary Table 1).

**Table 1.** Summary of the dataset.

| **Sample** | **Location** | **Type** | **Instrument** | **Clean Reads** | **Clean Bases** | **%GC** | **>Q20** | **>Q30** |
| --- | --- | --- | --- | --- | --- | --- | --- | --- |
| A | China | PE100 | BGISEQ-500 | 607,199,826 | 91,079,114,100 | 41 | 96.14 | 90.71 |
| A | South Africa | PE100 | MGISEQ-2000 | 607,871,944 | 91,079,892,900 | 41 | 96.44 | 91.47 |
| B | South Africa | PE100 | MGISEQ-2000 | 606,752,354 | 90,746,832,900 | 40 | 96.97 | 90.01 |
| C | South Africa | PE100 | MGISEQ-2000 | 608,824,594 | 91,180,791,600 | 41 | 97.01 | 92.55 |
| D | South Africa | PE100 | MGISEQ-2000 | 606,900,390 | 90,205,943,600 | 41 | 97.41 | 90.59 |
| E | South Africa | PE100 | MGISEQ-2000 | 608,317,592 | 90,770,106,600 | 41 | 95.67 | 90.97 |
| F | South Africa | PE100 | MGISEQ-2000 | 608,653,994 | 90,899,675,100 | 41 | 97.15 | 90.48 |

**Figure 1.**
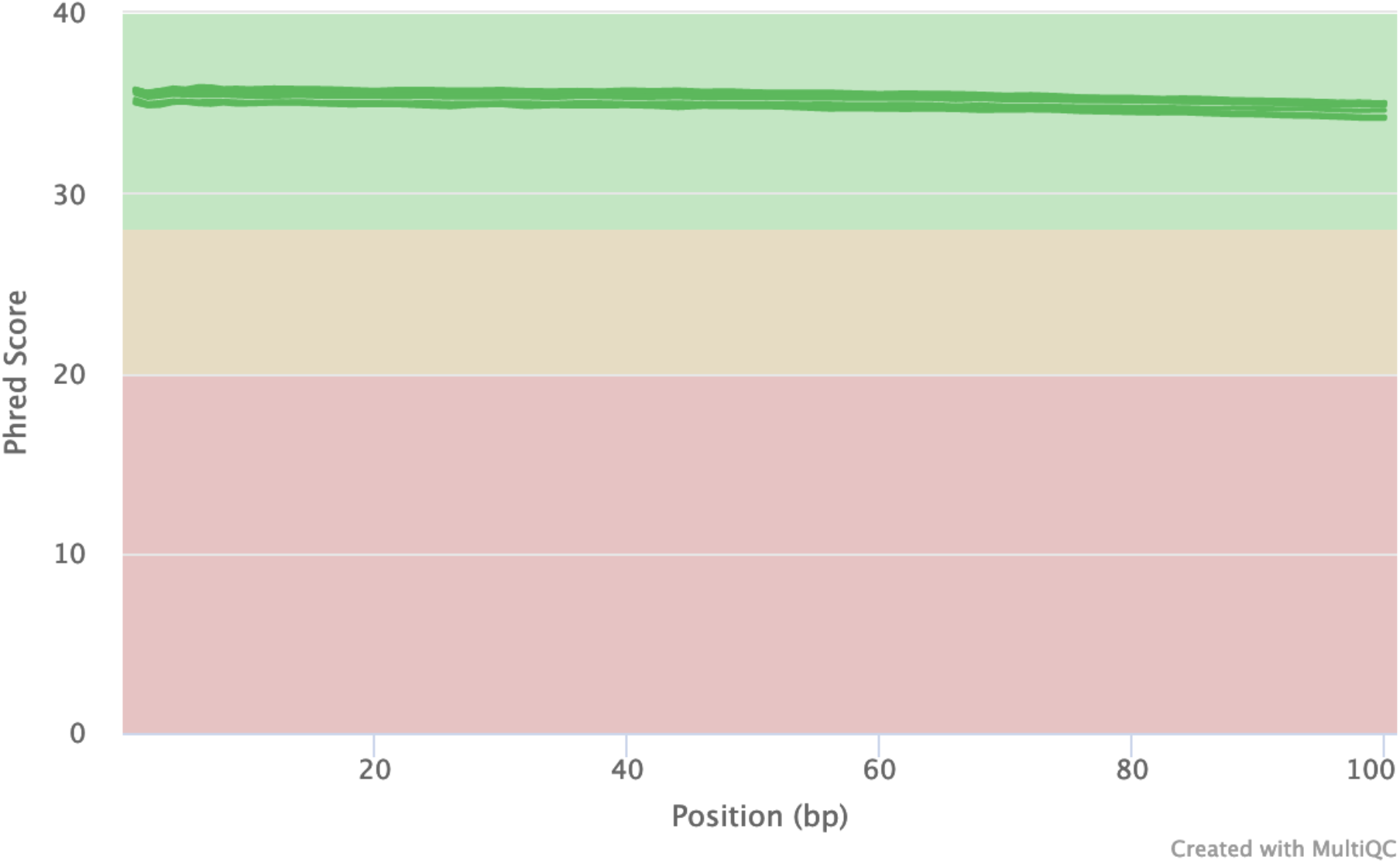
FastQC mean quality scores. FastQC quality scores for all seven samples were obtained. The higher the phred score, the better the base call. For all seven samples, bases for all samples were considered high quality (green). In addition, the quality scores remained consistent across the entire read length.

**Figure 2.**
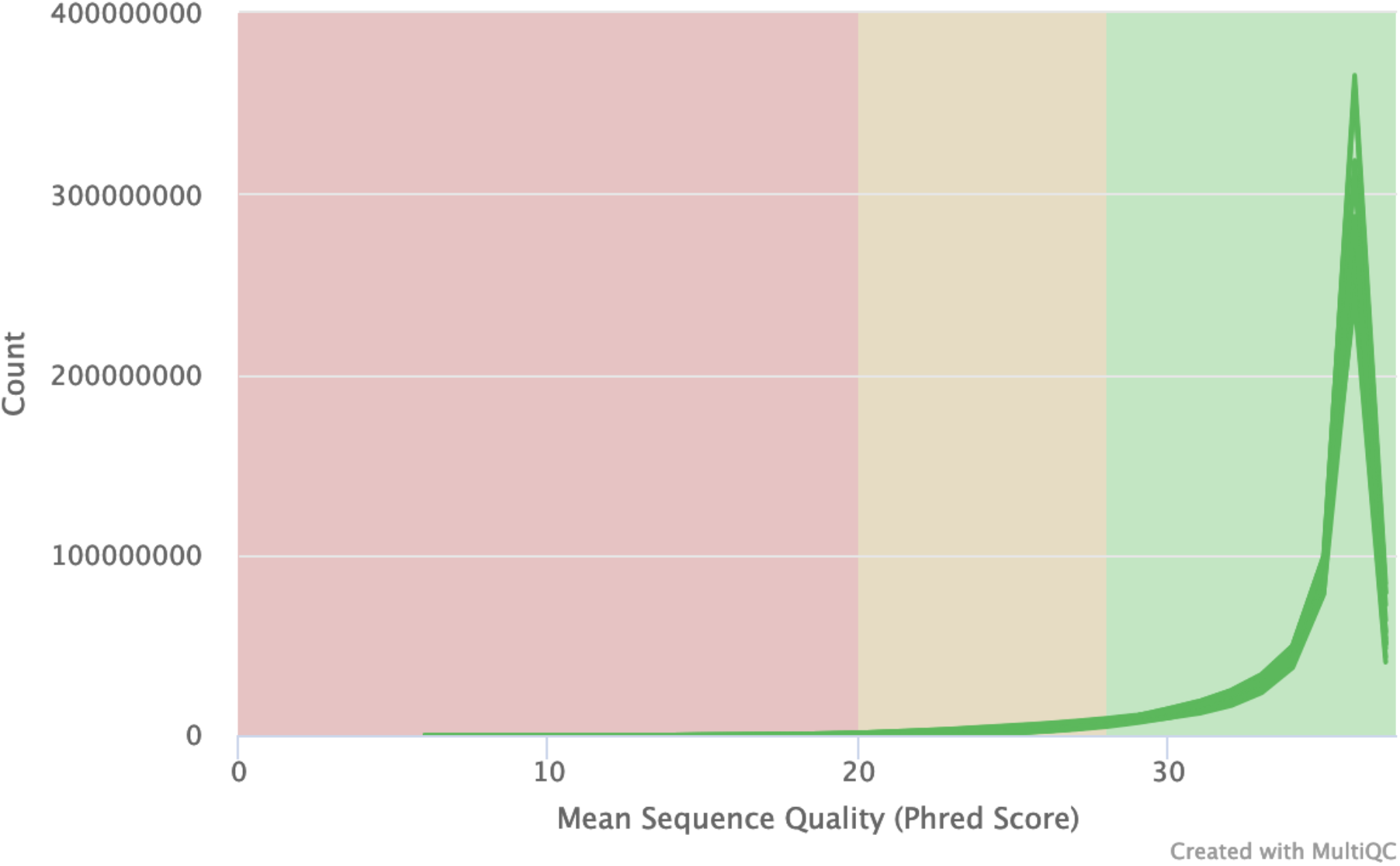
FastQC per sequence quality scores for all seven samples sequenced. All samples had universally high-quality scores.

**Figure 3.**
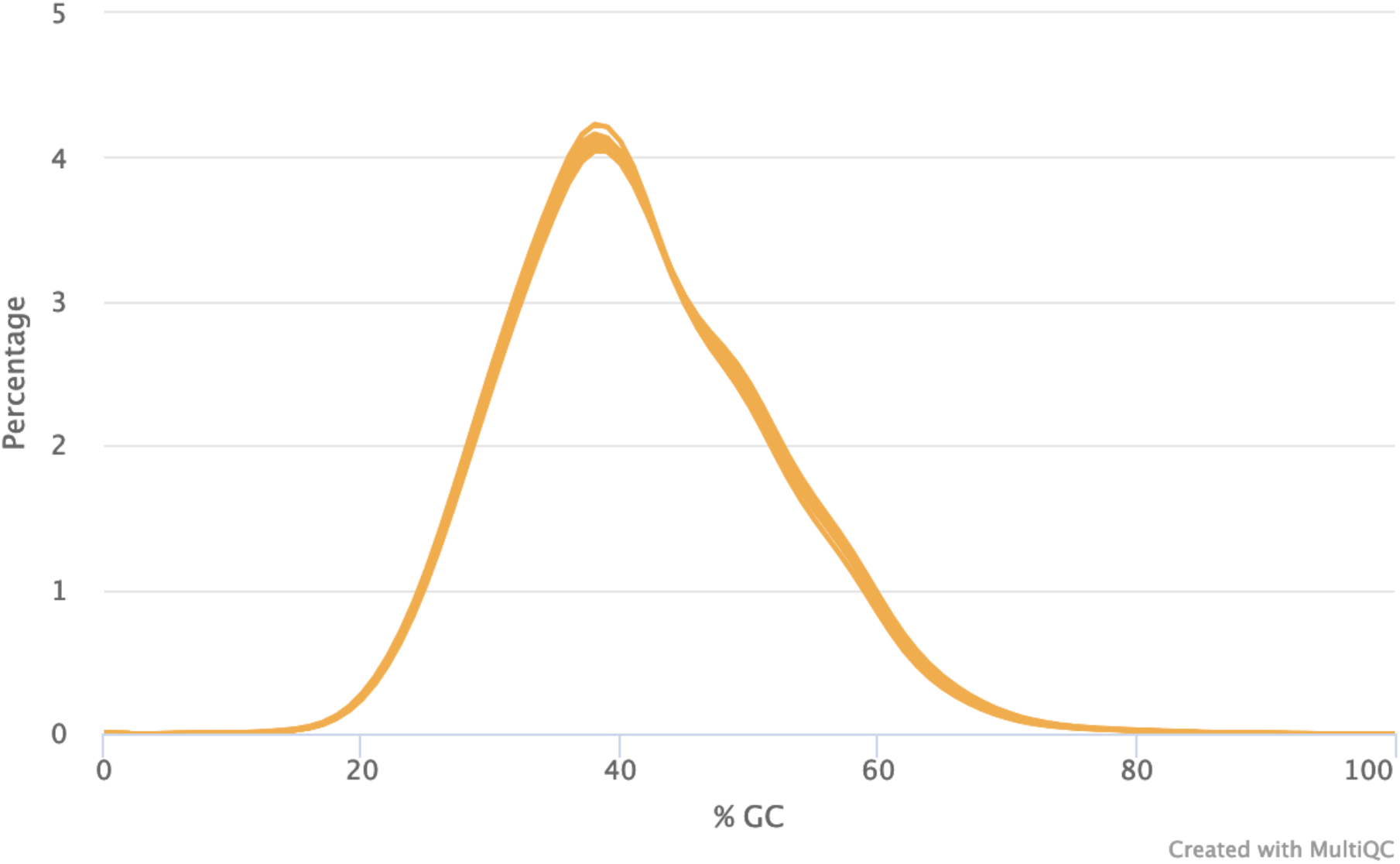
FastQC per sequence GC content. The GC content for all seven samples is normally distributed.

**Figure 4.**
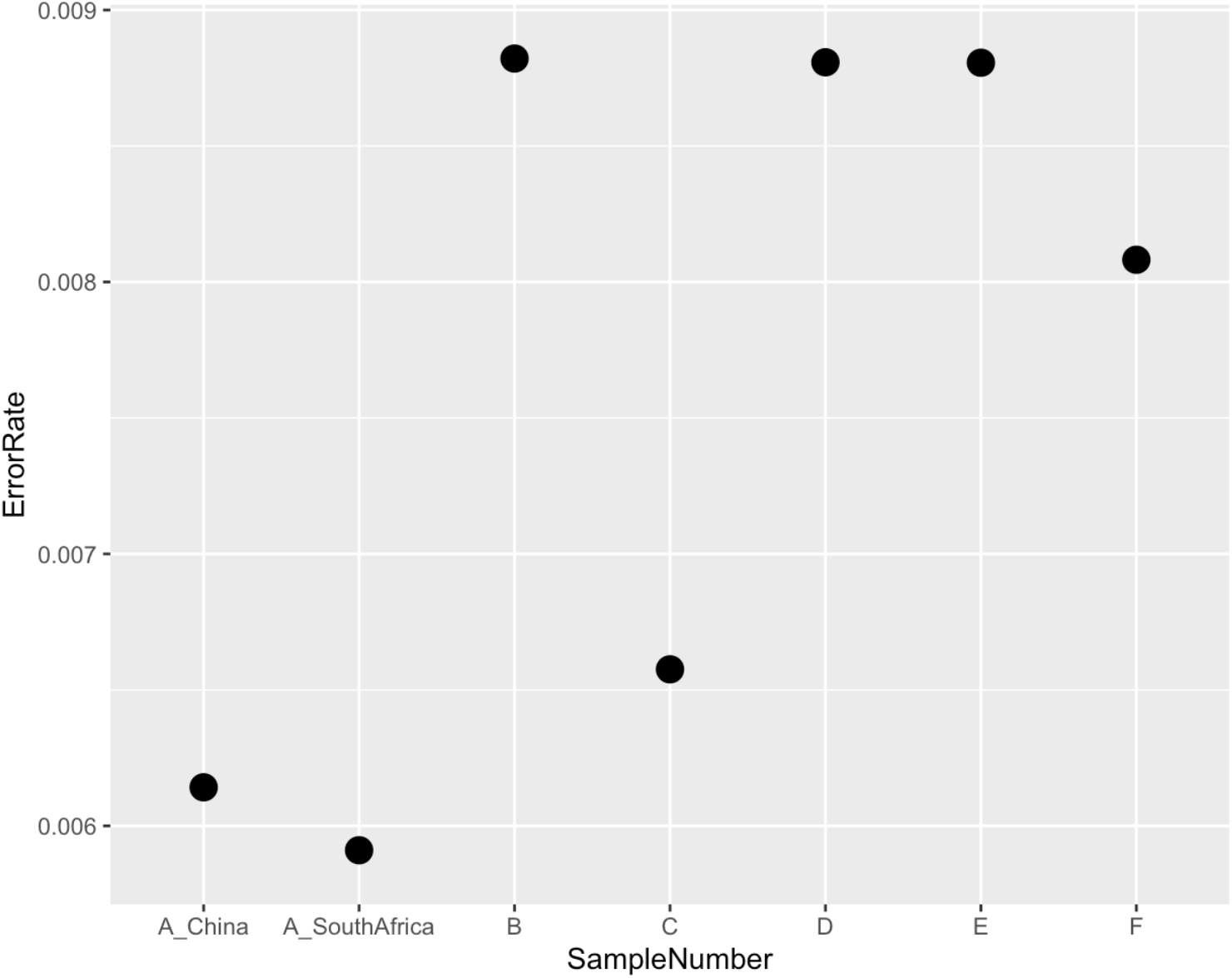
Error rates of each of the samples sequenced. The error rate is calculated using mismatches per base mapped.

**Figure 5.**
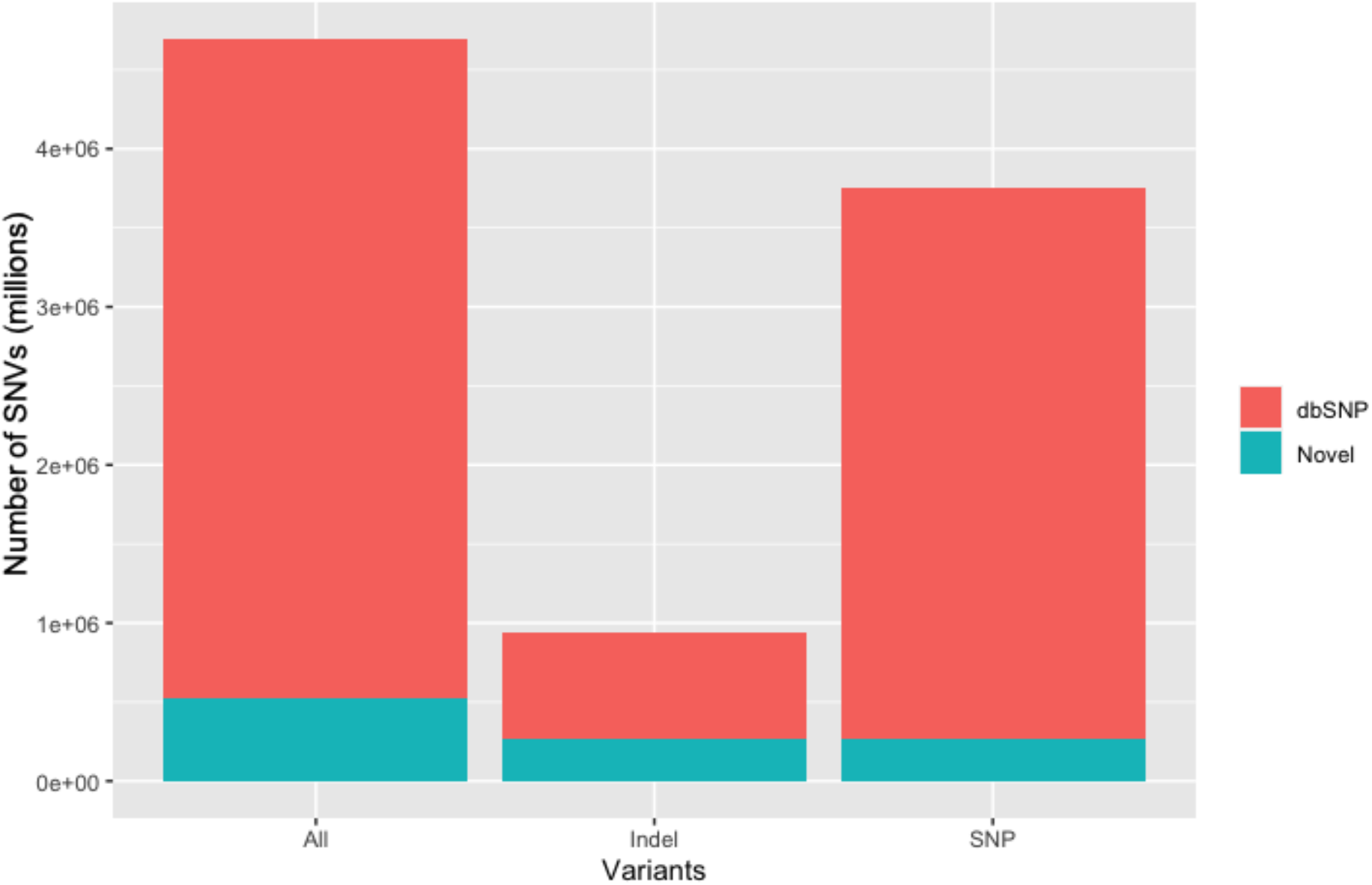
Average number of single nucleotide variants (SNVs) in all of the samples.

### Comparison of data obtained from MGISEQ-2000 and BGISEQ-500

Sample A was sequenced at both the SAMRC Genomics Centre in Cape Town, South Africa as well as at the BGI in China for comparative purposes. The overall results are illustrated in Table 2. There is a 99.91% similarity in the mapping rates of the two different platforms. The total read length for both platforms was 100bp and was maintained for both platforms with a coverage of 36.41X and 36.32X respectively (Figure 6). However, the sample sequenced on the MGISEQ-2000 platform had lower duplication (9.73% vs 10.12%) and overall mismatch rates (0.47% vs 0.51%) (Table 2). In addition, the overall number of clean reads was marginally higher on the MGISEQ-2000 with 778,800 more clean reads than those produced on the BGISEQ-500 for the same sample (Table 2).

**Table 2.** Comparative analysis of MGISEQ-2000 in South Africa with that of BGISEQ-500 at BGI.

|  | **Sample A in China** | **Sample A in South Africa** | **Percentage similarity** |
| --- | --- | --- | --- |
| **Instrument** | BGISEQ-500 | MGISEQ-2000 |  |
| **Minimum coverage 4X** | 99.02% | 99.88% | 99.14 |
| **Minimum coverage 10X** | 98.37% | 98.68% | 99.69 |
| **Minimum coverage 20X** | 96.62% | 96.96% | 99.65 |
| **Average depth** | 36.32 | 36.41 | 99.75 |
| **Clean reads** | 607,199,286 | 607,871,944 | 99.89 |
| **Clean bases** | 91,079,114,100 | 91,079,892,900 | 99.99 |
| **Identified bases** | 2,974,798,318 | 2,970,768,782 | 99.86 |
| **GRCh38.p13 length** | 3,272,116,950 | 3,272,116,950 | - |
| **Mapping rate** | 99.07% | 99.16% | 99.91 |
| **Duplication rate** | 10.12% | 9.73% | 96.15 |
| **Mismatch rate** | 0.51% | 0.47% | 92.16 |

**Figure 6.**
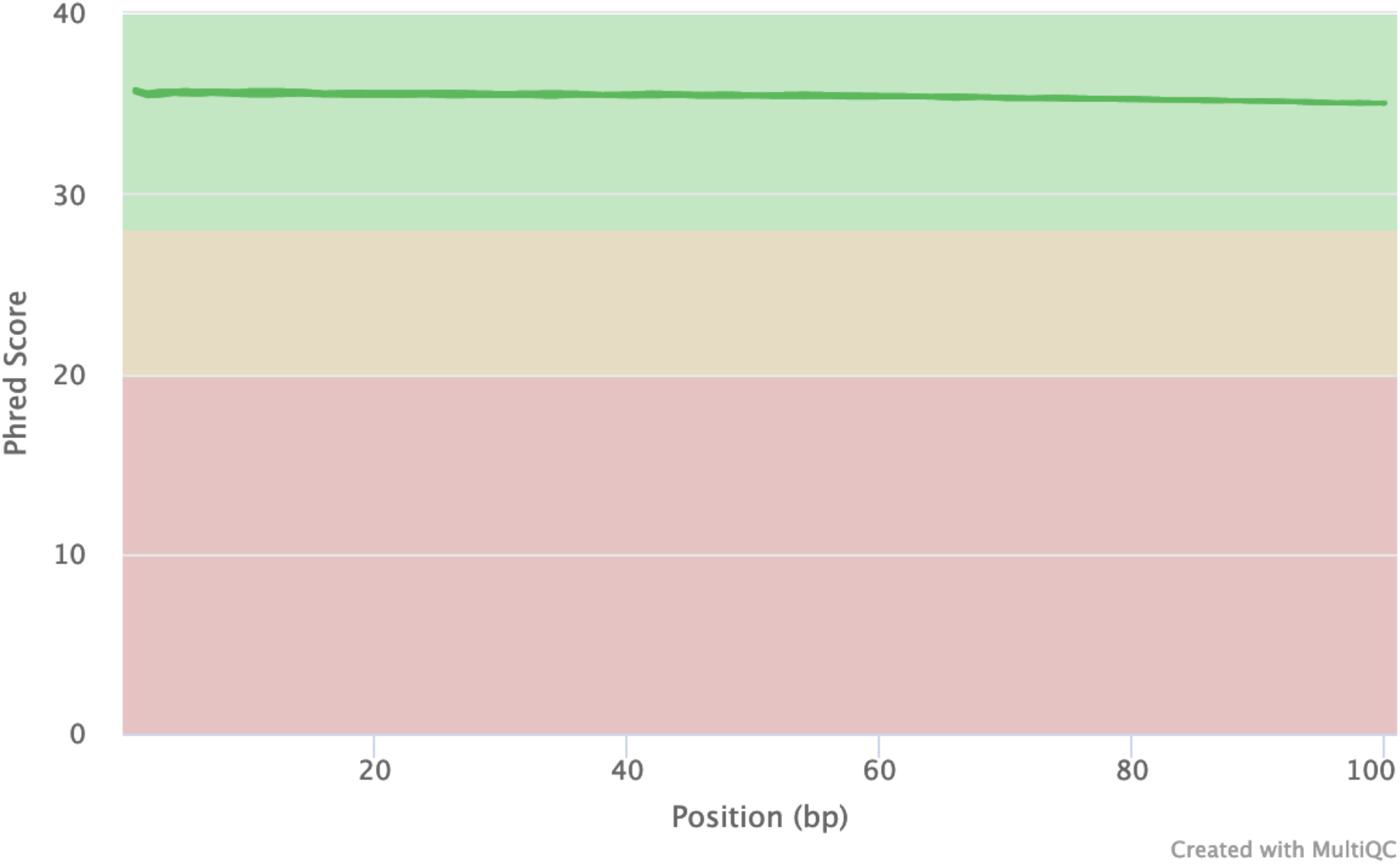
FastQC mean quality scores for Sample A. FastQC quality scores for Sample A were obtained. The quality scores are representative of those obtained from both the MGISEQ-2000 in South Africa and the BGISEQ-500 in China.

## Discussion

The data generated for this study is the first report of high-coverage WGS performed and analysed in South Africa at the SAMRC Genomics Centre. Data produced at the SAMRC Genomics Centre is of high quality with an excess of 30X coverage across the entire read length of 100bp, with coverage distribution almost identical across all samples. The data generated in South Africa is comparable to that produced at the BGI in China.

The Genome in a Bottle Consortium provides reference genomes for benchmarking, but we opted to use a South African DNA sample for comparison, as the same platforms and not manufacturers were compared ^14^. The data produced demonstrated the overall similarity of two different platforms designed and utilised by the BGI. The overall sequencing quality was higher on the MGISEQ-2000 when compared to the BGISEQ-500, with more clean bases and clean reads produced. The sequencing technology implemented on each platform is the same - with the generation of a DNA Nanoball (DNB) and the cPAS method, where an oligonucleotide probe is added and attaches in combination to specific sites within the DNB ^4,5^. Differences between the platforms may become clearer if longer read lengths are used (PE150) as read quality decreases over the entire read length. The technology on the MGISEQ-2000 is more advanced and the platform is able to produce up to 1500M - 1800M effective reads per flow cell (approximately 720GB data per single run) compared to the BGISEQ-500, which can only produce a maximum of 1300M effective reads, which equates to 520GB per run ^15^. This analysis demonstrated that the two instruments provide similar sequencing quality. The decrease in duplication rate is important as lower levels of duplication indicate high levels of coverage for a target sequence, whereas high levels indicate an enrichment bias.

In addition, our findings complement that of the SAHGP, which conducted deep sequencing (~50X) of 24 individual whole genomes ^8^. The SAHGP was the first high-coverage WGS study analysed and interpreted in South Africa with full funding from the South African government. The SAHGP had a higher coverage (47.66 vs 36.41) but the same read length of 100bp paired end was used for both projects. In 2017, the SAHGP detected 815,404 novel variants in 24 individuals - defined as absent from dbSNP build 142 ^16^, 1KGP ^17^ and the African Genome Variation Project (AGVP) ^18^. Our study detected 456,583 novel variants (230,432 SNPs and 226,151 indels) in only six individuals, demonstrating the genetic diversity present in South African individuals. The genomes in our study were also aligned to a newer reference than that of the SAHGP. While the present study did not make use of deep sequencing, the overall number of clean reads obtained was higher than that of the SAHGP, with an average of 9,085,165,257 clean reads across all samples. The current study was not only analysed and funded locally but was also completed using a WGS platform installed on the African continent and operated by South Africans.

The SAMRC Genomics Centre provides African researchers with the platform to better understand the factors which impact the individual and improve the response to disease. In addition, the local, state-of-the-art infrastructure enables researchers to explore avenues of research which may have been restricted due to limited infrastructure or budget constraints.

## Methods

### Study participants and ethics approval

Samples from six South African participants were available for sequencing as part of the platform installation. Participants were recruited from three sites as part of independent research projects. These studies were approved by the Health Research Ethics Committee of Stellenbosch University (Study no. N09/08/224 and Study no. N13/05/075(A)) and the Human Research Ethics Committee of the University of the Witwatersrand (Study no. M170585). Samples A and B were collected from two related individuals for a study investigating primary immunodeficiencies, and sample C was part of an HIV study. Samples D, E and F were recruited as part of a data sharing study of complex cases to determine whether WGS confirms the detection of a rare beta-isoform *TP53* variant [g.7576633A>G; NM_001126114.2: *TP53* c.1018A>G (p.N340D)] ^19^ as the most likely cause of Li Fraumeni-like syndrome previously detected using a pathology-supported genetic testing framework as previously described by van der Merwe et al. (2017) ^20^ In addition, one sample (Sample A) was previously subjected to WGS at the BGI using the BGISEQ-500.

### DNA extraction and quality assessment

Genomic DNA (gDNA) was extracted by three provider sites following their preferred standard protocols. Upon receipt of the DNA samples at the SAMRC Genomics Centre, a Quality Control (QC) Standard Operating Procedure (SOP) was followed. Genomic DNA samples were quantified with fluorometry using the Qubit 4.0 Fluorometer (Thermo Fisher Scientific, Waltham, MA, USA) and the Qubit dsDNA HS Assay kit according to the manufacturer’s instructions. Spectrophotometry was performed using the NanoDrop™ One Spectrophotometer (Thermo Fisher Scientific, Waltham, MA, USA) to determine the purity of the gDNA samples (A260/A280 and A260/230 ratio). As an additional assessment of the intactness, or the extent of possible degradation of the gDNA, all samples were resolved on an ethidium bromide pre-stained 1% agarose gel. Gel electrophoresis was carried out at 120 V in 1X SB buffer. All samples that met the QC criteria of a 260/280 ratio within the range of 1.8 and 2.2, a 260/230 ratio of above 1.7, with a gDNA yield greater than 500ng, and a high integrity (high molecular weight with intact dsDNA and no secondary bands on an agarose gel), underwent library construction.

### Library construction and whole genome sequencing

The gDNA samples (1000ng) were subjected to physical shearing with the M220 Focused-ultrasonicator (Covaris, Woburn, MA, USA), followed by magnetic bead-based size selection using MGIEasy DNA Clean Beads (MGI, Shenzhen, China) prior to proceeding with library construction. Library preparation was performed with 50ng of fragmented DNA for each sample using the MGIEasy Universal DNA Library Prep Kit (MGI, Shenzhen, China), according to the manufacturer’s instructions. Briefly, each sample was subjected to an End-repair and A-tailing (ERAT) reaction, using the appropriate volumes of ERAT Buffer and ERAT Enzyme mix. The end-repaired products were ligated to MGIEasy DNA Adapters as per the manufacturer’s guidelines. Adapter-ligated DNA was purified using MGIEasy DNA Clean Beads and amplified using the MiniAmp™ Thermal Cycler (Thermo Fisher Scientific, Waltham, MA, USA). PCR products were purified as previously described and quantified with fluorometry using the Qubit dsDNA HS Assay kit according to the manufacturer’s instructions. Additionally, the fragment size distribution of purified PCR products was assessed using gel electrophoresis. Single-stranded, circular DNA libraries were generated from 1pmol of purified PCR product for each sample, followed by purification and quantification with MGIEasy DNA Clean Beads and the ssDNA HS Assay kit (Qubit), respectively. The MGILD-200 automatic loader was used to load sample libraries onto the MGISEQ-2000 FCL flow cells. Massively parallel sequencing was performed using DNA nanoball-based technology on the MGISEQ-2000 (BGI, Shenzhen China) with the appropriate reagents supplied in the MGISeq-2000RS High-Throughput Sequencing Kit. A paired-end sequencing strategy was employed, with a read length of 100bp (PE100).

### Sequencing quality check, mapping, and data analysis

All data sets were processed locally using South African computational infrastructure. Raw datasets were transferred to the Centre for High Performance Computing’s Lengau cluster, where all downstream analyses were conducted. FastQC (version 0.11.9) was used to check the sequence quality, and Q20/Q30 ratios were calculated using q30, a freely available Python script ^21^. Raw data sets were pre-processed using Trimmomatic ^22^ which included the removal of adapter sequences, low quality reads as well as very short reads (<20bp). Genome Analysis Toolkit (GATK) version 4.0 framework was used for all downstream processing of the data ^23^. Burrows-Wheeler Aligner (BWA)-MEM (version 0.7.17), with default parameters, was used to align all “cleaned” sequencing reads to the human reference genome GRCh38p13 ^24^. The quality of the aligned reads was assessed using SAMtools (version 1.9) ^13^. Duplicate reads were removed using Picard ^25^, followed by base quality score recalibration (BQRS) using the protocol provided by Genome Analysis Toolkit (GATK) ^26^. Variants were called using HaplotypeCaller ^27^ producing a variant called format (VCF) file. Following VCF file generation, variants were annotated using ANNOVAR software using the database version (2019Jun17) ^28^. Variants were classified as novel if they were absent from gnomAD ^29^, dbSNP (build 153) ^16^ and the 1000 Genomes Project (1KGP) ^17^. The novel germline *TP53* variant c.1018A>G (p.N340D) previously detected in sporadic hepatocellular carcinoma and endometrial cancer ^30^ served as an internal control following WGS data transfer.

## Data Availability

All whole genome sequencing data were aligned to human reference genome GRCh38 from the Genome Reference Consortium Human Build 38 patch release 13 (https://www.ncbi.nlm.nih.gov/assembly/GCF_000001405.39). The datasets generated and analysed during the current study are not publicly available as participants did not consent to this, but are available from CK (samples A-B), CTT (sample C) and MK (samples D-F) on reasonable request.

## Acknowledgements

We thank the Centre for High Performance Computing (https://www.chpc.ac.za/), Werner Janse van Rensburg and Inus Scheepers for providing access to computational infrastructure, as well as additional technical support. We would also like to thank Ms Anel Sparks for helping with the generation of images presented in this manuscript.

## Author contributions

BG, MM, RG, RiM, TJ, SG and CK conceptualised the study. All authors provided substantial intellectual input into the manuscript. MK, ME and CTT provided DNA samples for sequencing after obtaining ethics approval and informed consent from patients. RiM and RoM designed and established the laboratory. CK and SG assisted in laboratory equipment procurement. HL trained and supervised TJ and SG who performed the sequencing. PC and CD were responsible for data management. BG and AP processed the data. BG analysed the data under the supervision of MM and CK. BG, MM, TJ, SG and CK drafted the manuscript. GG is the president of the SAMRC and RiM, GG and GL secured funding for the Genomics Centre. BG, MM and RJW secured funding for sequencing at BGI. JL and RMW contributed scientific discussion. All authors approved the final manuscript as submitted and agree to be accountable for all aspects of the work.

