## Supplementary Figures and Tables for "Human Whole Genome Sequencing in South Africa"

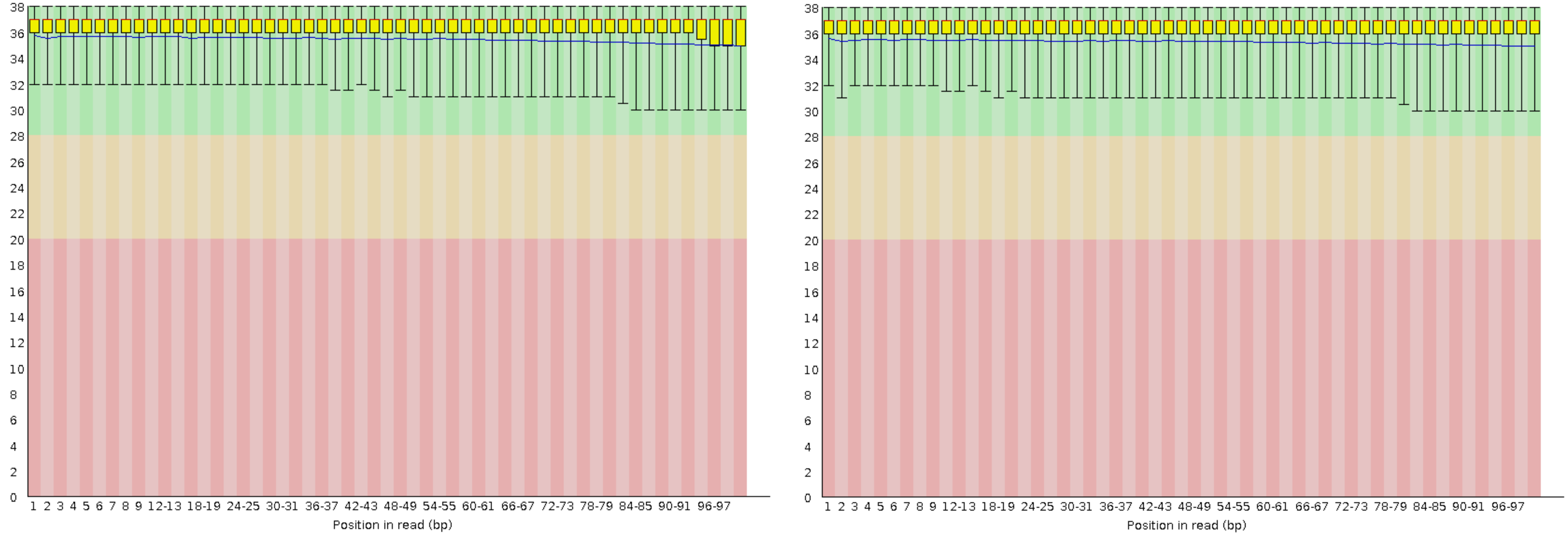


**A** **B**

**Supplementary Figure 1A and B.** Distribution of nucleotide quality parameters across forward (A) and reverse (B) reads for sample A sequenced at the Beijing Genomics Institute, China.

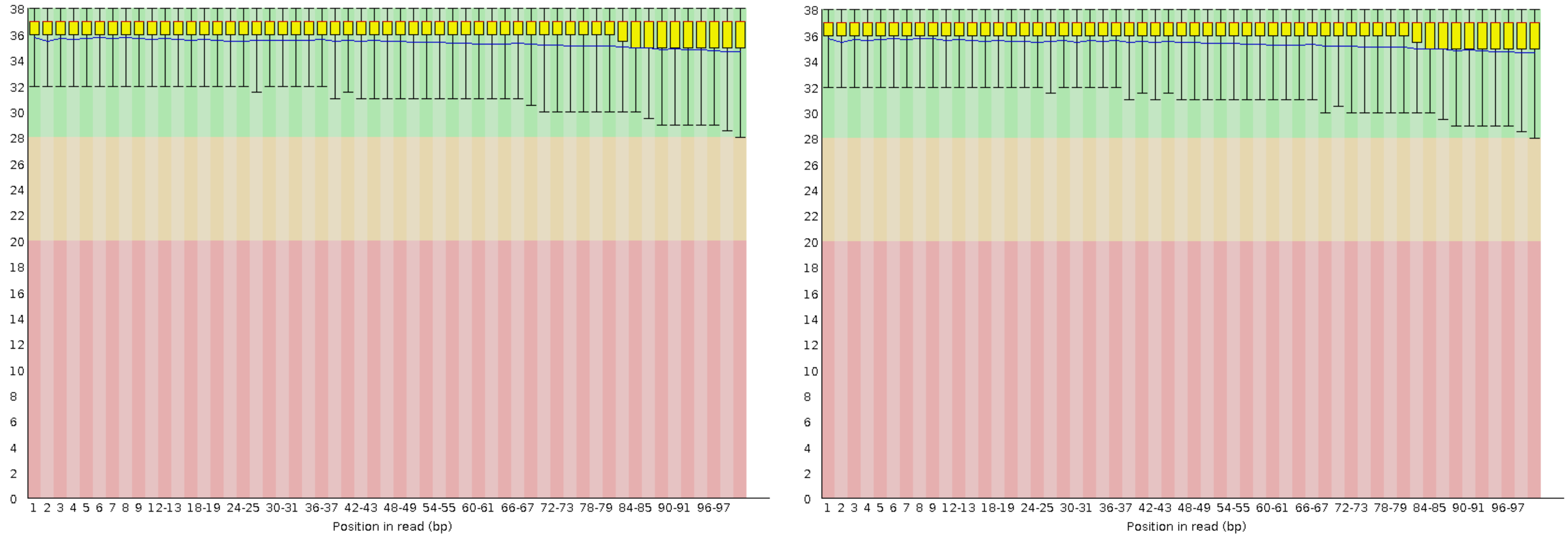


**A** **B**

**Supplementary Figure 2A and B.** Distribution of nucleotide quality parameters across forward (A) and reverse (B) reads for sample A sequenced at the Genomics Centre, South Africa.

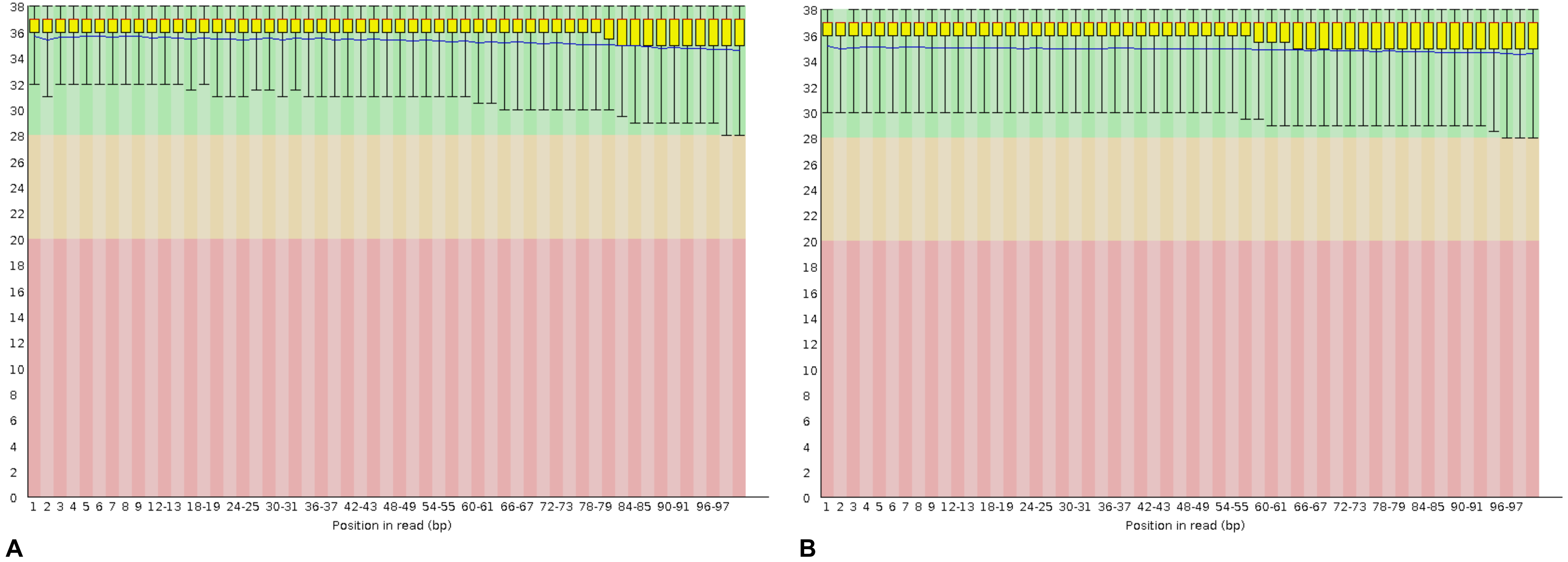


**Supplementary Figure 3A and B.** Distribution of nucleotide quality parameters across forward (A) and reverse (B) reads for sample B sequenced at the Genomics Centre, South Africa.

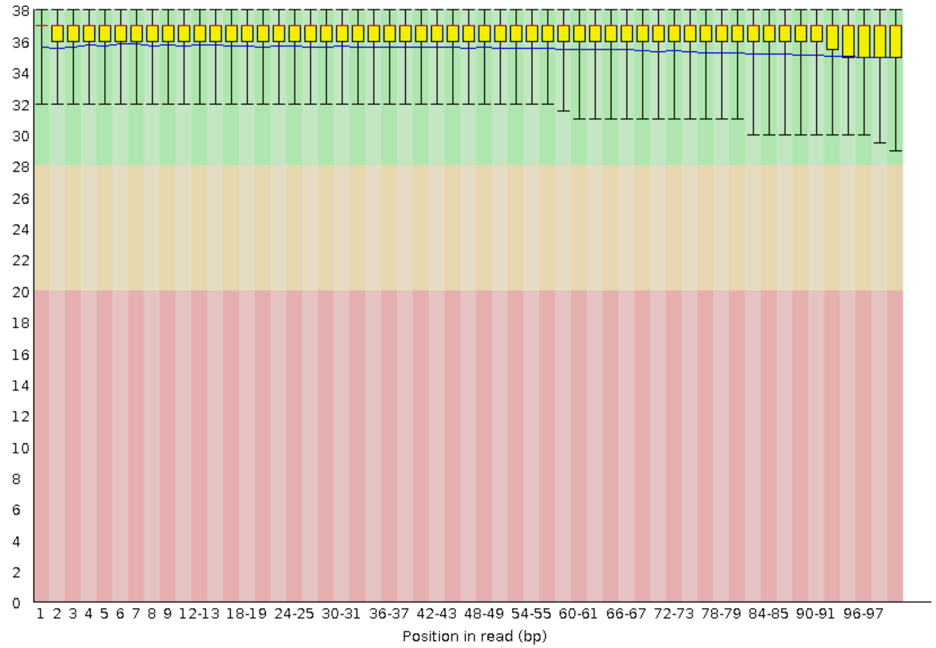


**A**

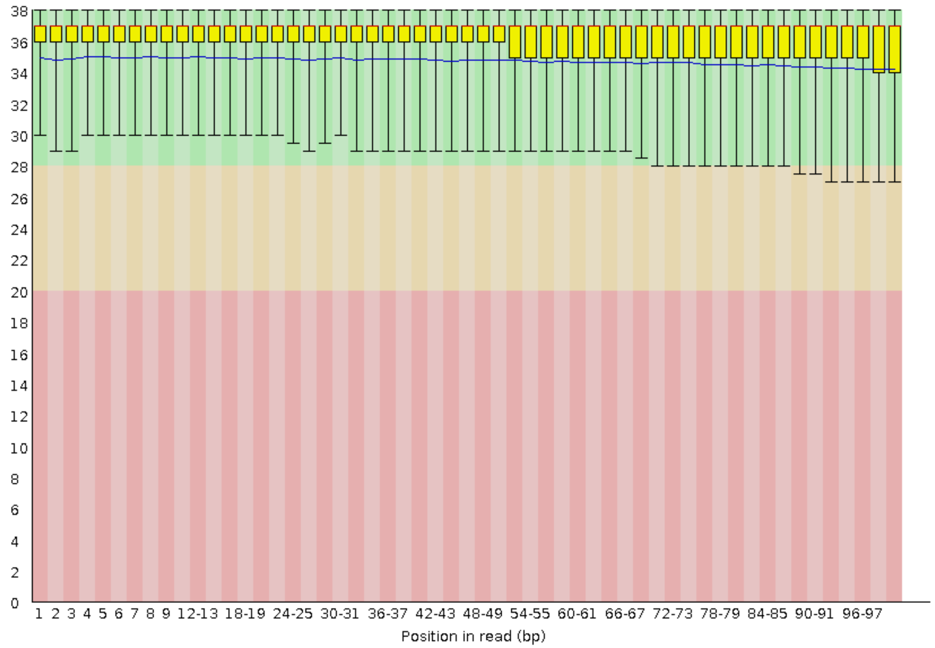


**B**

**Supplementary Figure 4A and B.** Distribution of nucleotide quality parameters across forward (A) and reverse (B) reads for sample C sequenced at the Genomics Centre, South Africa.

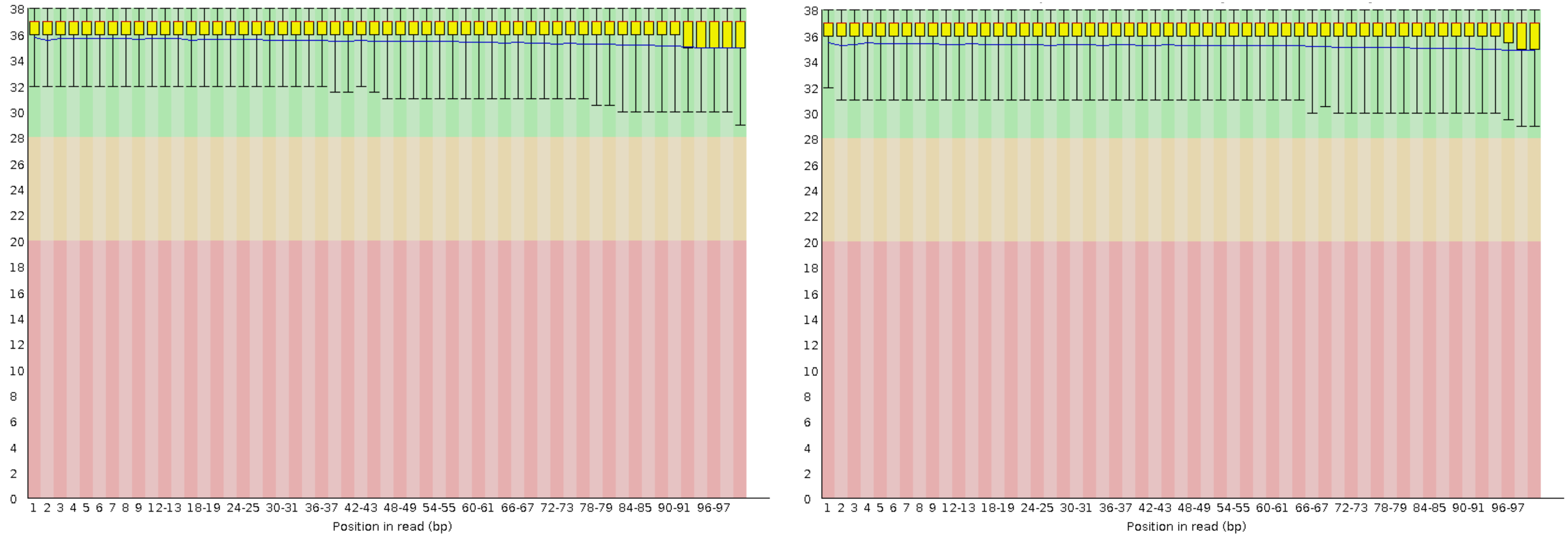


**A** **B**

**Supplementary Figure 5A and B.** Distribution of nucleotide quality parameters across forward (A) and reverse (B) reads for sample D sequenced at the Genomics Centre, South Africa.

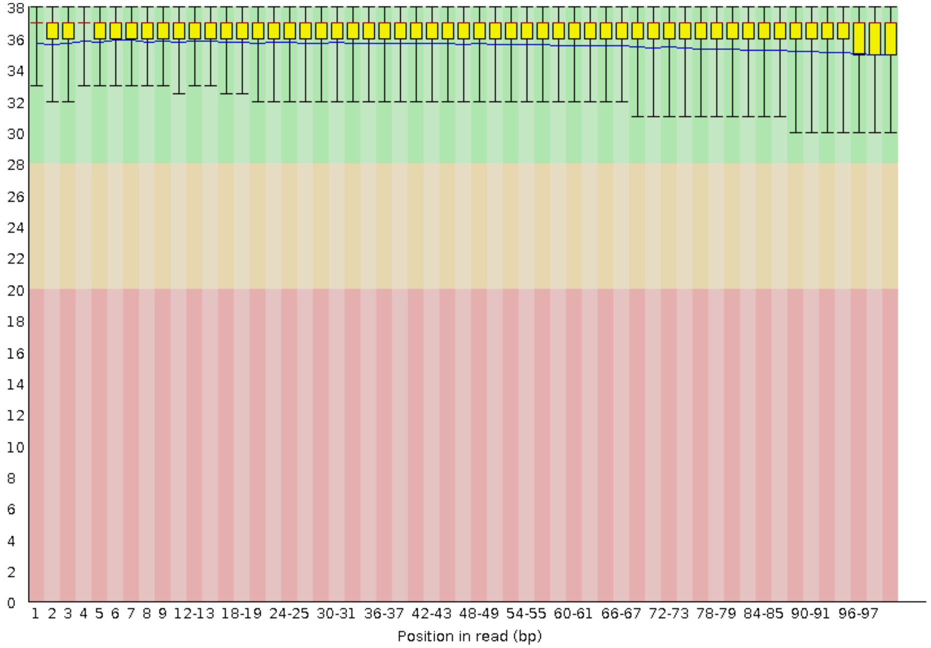


**A**

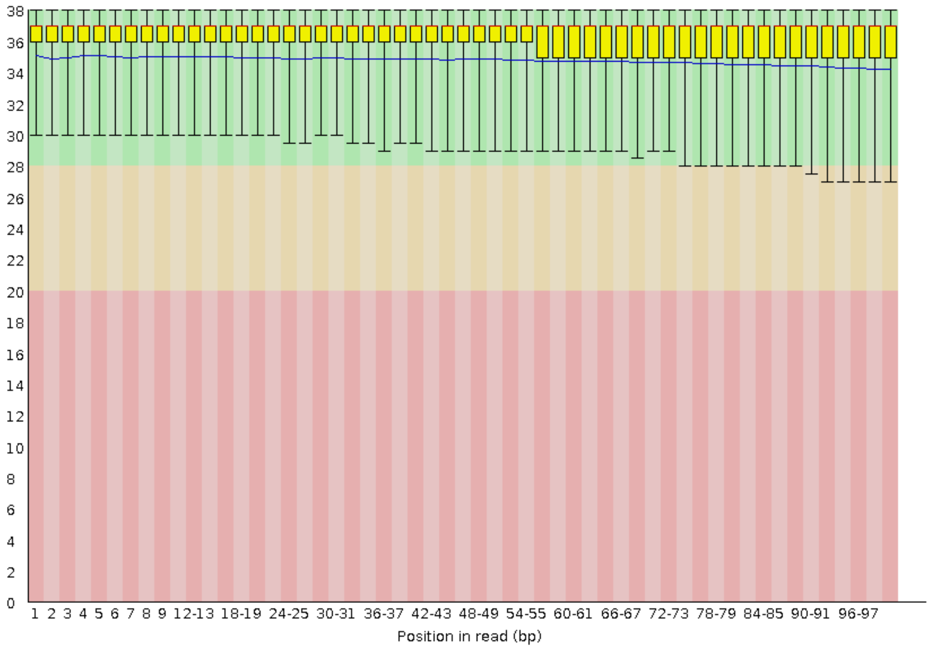


**B**

**Supplementary Figure 6A and B.** Distribution of nucleotide quality parameters across forward (A) and reverse (B) reads for sample E sequenced at the Genomics Centre, South Africa.

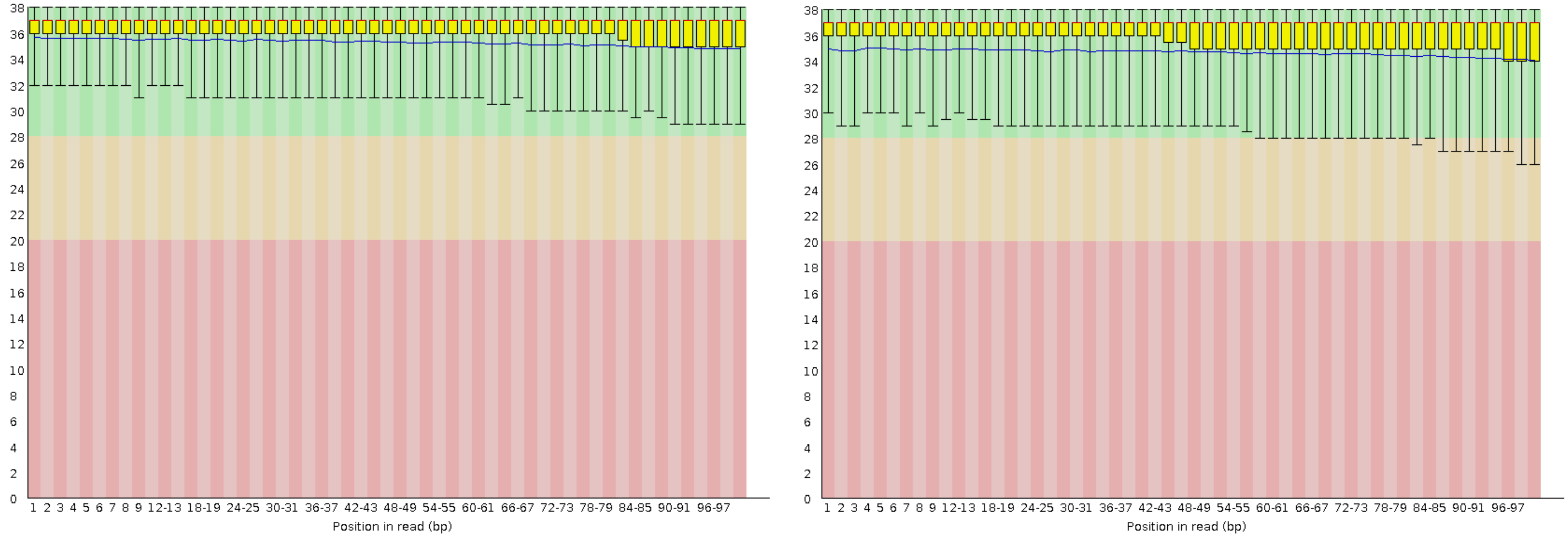


**A** **B**

**Supplementary Figure 7A and B.** Distribution of nucleotide quality parameters across forward (A) and reverse (B) reads for sample F sequenced at the Genomics Centre, South Africa.

.

**Supplementary Table 1: Summary of all variants identified in all 7 samples.**

| **Sample** | **A in China** | **A in South Africa** | **B** | **C** | **D** | **E** | **F** |
|---|---|---|---|---|---|---|---|
| **Instrument** | BGISEQ-500 | MGISEQ-2000 | MGISEQ-2000 | MGISEQ-2000 | MGISEQ-2000 | MGISEQ-2000 | MGISEQ-2000 |
| **Total number of variants** | **4,841,151** | **4,847,679** | **4,753,897** | **4,781,930** | **4,485,241** | **4,563,634** | **4,592,587** |
| **SNPs** | 3,907,017 | 3,913,468 | 3,815,771 | 3,842,348 | 3,531,385 | 3,623,269 | 3,636,764 |
| **Found in dbSNP** | 3,617,507 | 3,615,260 | 3,549,049 | 3,579,916 | 3,290,191 | 3,358,408 | 3,386,191 |
| **Novel** | 289,510 | 298,208 | 266,722 | 262,432 | 241,194 | 264,861 | 250,573 |
| **Homozygous** | 1,683,211 | 1,680,224 | 1,638,110 | 1,677,224 | 1,478,591 | 1,561,991 | 1,555,808 |
| **Heterozygous** | 2,223,806 | 2,233,244 | 2,177,661 | 2,165,124 | 2,052,794 | 2,061,278 | 2,080,956 |
| **Indels** | 934,134 | 928,211 | 938,126 | 939,582 | 953,856 | 940,365 | 958,823 |
| **Found in dbSNP** | 679,302 | 681,493 | 685,864 | 668,325 | 686,586 | 668,694 | 680,573 |
| **Novel** | 254,832 | 246,718 | 252,262 | 271,257 | 258,495 | 271,671 | 278,250 |
| **Homozygous** | 134,809 | 138,579 | 140,062 | 139,152 | 134,875 | 133,438 | 135,769 |
| **Heterozygous** | 799,342 | 789,632 | 798,064 | 800,430 | 818,981 | 806,927 | 823,054 |
